# Bacterial type II topoisomerases cleave DNA in a species-specific manner

**DOI:** 10.1101/2025.07.28.667256

**Authors:** Ian L. Morgan, Jeffrey Y. Jian, Neil Osheroff, Keir C. Neuman

**Affiliations:** National Heart, Lung, and Blood Institute, National Institutes of Health, Bethesda, MD 20892, USA; Department of Biochemistry, Vanderbilt University School of Medicine, Nashville, TN 37232, USA; Department of Medicine, Vanderbilt University School of Medicine, Nashville, TN 37232, USA

## Abstract

The type II topoisomerases, gyrase and topoisomerase IV, are essential enzymes in nearly all bacteria and are the targets of fluoroquinolones, which are some of the most widely prescribed broad-spectrum antibacterials in clinical use. As part of their catalytic cycle, gyrase and topoisomerase IV transiently cleave DNA in a sequence-dependent manner. However, it is unclear whether this sequence-dependence is species-specific. Therefore, using our recently developed SHAN-seq method, we mapped and compared cleavage sites for type II topoisomerases from three different pathogenic bacterial species, *Escherichia coli, Bacillus anthracis*, and *Mycobacterium tuberculosis* in the presence of the fluoroquinolone, ciprofloxacin. We found that the enzymes have substantially different DNA cleavage specificities that vary between gyrase and topoisomerase IV, across species, with supercoil chirality, and in response to ciprofloxacin. Our results demonstrate that bacterial species fine-tune the DNA cleavage specificity of their type II topoisomerases. This finding suggests that cleavage specificity may play important physiological roles and, in turn, may affect the susceptibility of bacteria to fluoroquinolone antibacterials.

## Introduction

The type II topoisomerases, gyrase and topoisomerase IV, are essential enzymes in nearly all bacteria and the molecular targets of fluoroquinolones, which are some of the most widely prescribed broad-spectrum antibacterials in clinical use (1–3). These enzymes are responsible for resolving the DNA entanglements (e.g., supercoils and catenanes) that arise during transcription and replication (4) and that can impede gene expression and chromosome segregation. To resolve these entanglements, gyrase and topoisomerase IV use ATP to capture and pass one double-stranded DNA segment through another (1, 2). As part of this process, the enzymes transiently cleave one of the DNA segments and form a protein-DNA intermediate called a cleavage complex. Since DNA cleavage can cause genome instabilities, such as DNA damage, mutations, and translocations, bacteria need to carefully regulate the type II topoisomerases DNA cleavage-religation equilibrium to ensure proper localization and activity (4, 5). Fluoroquinolones disrupt this equilibrium by preventing type II topoisomerase DNA religation, trapping the enzyme in a cleavage complex. Hence, the regulation of DNA cleavage mediated by type II topoisomerases is integral to both their physiological roles and their roles as the molecular targets of antibacterials.

Two major factors regulate the DNA cleavage activity of type II topoisomerases: DNA topology and sequence (6, 7). Type II topoisomerases respond to supercoiling, generating higher cleavage levels on supercoiled DNA than on relaxed DNA (7). This increased cleavage of supercoiled DNA is thought to help the enzymes resolve DNA entanglements (7). Notably, gyrase also recognizes supercoil chirality, maintaining higher cleavage levels on negatively supercoiled (−sc) DNA than on positively supercoiled (+sc) DNA (5-7). This supercoil-dependent difference in cleavage-religation equilibrium is thought to help avoid genomic instabilities associated with collisions between cleavage complexes and transcription complexes, which generate +sc downstream, and -sc upstream, of transcription (8, 9).

In addition to DNA topology, DNA sequence also influences the gyrase and topoisomerase IV DNA cleavage-religation equilibrium. Depending on the surrounding DNA sequence, some sites have higher type II topoisomerase DNA cleavage levels than others (10–14). In contrast to restriction enzymes, type II topoisomerases do not have a single well-defined recognition sequence (12–15). Instead, they cleave many DNA sites with different sequences, albeit with widely varying frequencies (15). The physiological implications of these sequence preferences are still emerging, but several ‘strong’ cleavage sites have been shown to have important biological roles. For example, the *Escherichia coli* (*Ec*) Mu bacteriophage contains a strong gyrase site that is essential for efficient recombination of the integrated viral genome (16–18). Yet, the factors that determine type II topoisomerase sequence preferences remain unclear.

The physical interactions between type II topoisomerases and DNA are thought to play an important role in their sequence preferences. Type II topoisomerases interact with DNA via their cleavage and C-terminal domains (CTDs) (19–21). The cleavage domains of gyrases and topoisomerase IV have a similar structure and sequence and bind approximately 20 base pairs (bp) surrounding the cleavage site (19, 20). The sequence identity of the bound DNA significantly influences the cleavage-religation equilibrium established by the enzyme (12, 13, 15, 22, 23). The CTDs have a similar beta-pinwheel architecture and a series of repeating Greek key motifs called ‘blades’ that interact with DNA but have some notable sequence and structural differences (24–26). The CTDs of gyrase have a conserved GyrA-box motif and six blades that can fully wrap up to 120-140 bp of DNA (21, 27). In contrast, the CTDs of topoisomerase IV interact with DNA, but lack the conserved GyrA-box and have variable numbers of blades, so they do not fully wrap the DNA (28). Whereas both enzymes can resolve a variety of DNA entanglements, the difference in their structures contributes to gyrase more efficiently resolving +sc, and topoisomerase IV more efficiently resolving catenanes.

Although gyrase and topoisomerase IV share substantial sequence and structural homology across bacterial species (26), previous work has shown that species-specific variations can affect their susceptibility to fluoroquinolones and catalytic activity (29). For example, within the cleavage domain, most gyrases and topoisomerases IV have two conserved residues, a serine and acidic residue, that stabilize fluoroquinolones via a water-metal ion bridge (2, 3, 30). Subtle variations in the orientation of these residues across bacterial species, such as in *Ec* and *Bacillus anthracis* (*Ba*), alters their response to fluoroquinolones (31–34). Additionally, a few bacterial species, such as *Mycobacterium tuberculosis* (*Mt*), have type II topoisomerases that lack one of these conserved residues, which accounts for their natural resistance to fluoroquinolones (35, 36). Some type II topoisomerases have species-specific insertions that influence their catalytic activity, such as a 170-amino acid insertion in *Ec* gyrase within the cleavage domain that has been implicated in DNA binding (37). The CTDs of topoisomerase IV also vary widely across species in their overall shape and number of blades, which influences the degree to which they interact with DNA (38). Furthermore, whereas gyrase CTDs have the same number of blades, they have species-specific C-terminal tails that regulate the extent to which gyrase can supercoil DNA (39). Each of these features could potentially influence the cleavage specificity of type II topoisomerases.

To determine the effects of species-specific variations on the type II topoisomerase DNA cleavage-religation equilibrium, we used our recently developed *in vitro* simple, high-accuracy end-sequencing method (SHAN-seq) (15) to map and compare the DNA cleavage specificities of gyrase and topoisomerase IV from three different pathogenic bacterial species, *Ec, Ba*, and *Mt*. We found that the enzymes have vastly different DNA cleavage specificities between gyrase and topoisomerase IV, across species, with supercoil chirality, and in response to the fluoroquinolone, ciprofloxacin. Our results suggest that bacteria fine-tune the specificity of DNA cleavage mediated by their gyrase and topoisomerase IV, which, in turn, may affect their susceptibility to fluoroquinolones.

## Methods

### Enzymes, DNA, and materials

We expressed and purified *Ba* gyrase subunits, GyrA and GyrB, and topoisomerase IV subunits, GrlA and GrlB, as well as *Mt* gyrase subunits, GyrA and GyrB as previously described (9, 31, 35). Unlike many bacterial species, *Mt* expresses gyrase but not topoisomerase IV (9, 35, 40). We expressed and purified human tyrosyl DNA phosphodiesterase-2 (TDP2) as previously described (15). We expressed and purified *Archaeoglobus fulgidus* reverse gyrase as previously described (41). We stored all enzymes at -80 °C.

We prepared -sc pBR322 plasmid DNA from *Ec* using a Plasmid Mega Kit (Qiagen) as described by the manufacturer. We prepared +sc pBR322 plasmid DNA by treating -sc DNA with recombinant *A. fulgidus* reverse gyrase as previously described (41). Briefly, reaction mixtures contained 35 nM (−)SC pBR322 DNA and 420 nM reverse gyrase in a total of 500 μl of 50 mM Tris-HCl (pH 8.0), 10 mM NaCl, 10 mM MgCl_2_, and 1 mM ATP. We incubated reactions at 95 °C for 10 min, stopped them by addition of 20 μl of 250 mM Na_2_EDTA, and cooled them on ice. We added 5 μl of Proteinase K (PK, Affymetrix, 8 mg/ml) to each reaction and incubated them at 45 °C for 30 min to digest the reverse gyrase. We extracted the DNA from each reaction with phenol:chloroform:isoamyl alcohol (25:24:1) and precipitated the DNA with 100% ice-cold ethanol. We resuspended the DNA in 50 μl of 5 mM Tris (pH 8.0), 0.5 mM EDTA (pH 8.0) and purified it using Bio-Spin P-30 gel columns (Bio-Rad, #7326231). We assessed the purity of DNA via the A260/A280 absorbance ratio. To ensure that differences between -sc and +sc DNA substrates were not influenced by temperature or other conditions used in the preparation protocol, we similarly treated -sc plasmid substrates in parallel with the omission of reverse gyrase.

We purchased analytical grade ciprofloxacin from LKT and stored it at 4°C as a 40 mM stock solution in 0.1 N NaOH.

### Bacterial type II topoisomerase cleavage assays

We performed DNA cleavage assays with *Ba* gyrase and topo IV and *Mt* gyrase as previously described (9, 31, 35). Due to the loss of DNA during the subsequent library preparation procedure, we included 5-fold more than the typical amount of DNA and enzyme. The *Ba* gyrase cleavage assays contained 250 nM enzyme (1:2 GyrA:GyrB ratio) and 50 nM -sc or +sc pBR322 DNA in 100 μl of 50 mM Tris-HCl (pH 7.5), 100 mM potassium glutamate (KGlu), 5 mM MgCl_2_, 5 mM dithiothreitol (DTT) and 50 μg/ml BSA. The *Ba* topo IV cleavage assays contained 750 nM enzyme (1:2 GrlA:GrlB ratio), 50 nM -sc or +sc pBR322 DNA in 100 μl of 40 mM Tris-HCl (pH 7.9), 50 mM NaCl, 5 mM MgCl_2_ and 2.5% glycerol (v/v). The *Ba* gyrase and topoisomerase IV cleavage assay reactions were incubated at 37 ºC for 30 or 10 min, respectively. The Mt gyrase cleavage assays contained 500 nM enzyme (1.5:1 GyrA:GyrB ratio) and 50 nM -sc or +sc pBR322 DNA in 100 μl of 10 mM Tris-HCl (pH 7.5), 40 mM KCl, 6 mM MgCl_2_, 0.1 mg/ml BSA, and 10% glycerol (v/v). The *Mt* gyrase cleavage assay reactions were incubated at 37 ºC for 10 min. All stated enzyme concentrations reflect that of the holoenzyme (A_2_B_2_). All cleavage reactions were carried out in the presence or absence of saturating ciprofloxacin (100 µM).

Following incubation of the cleavage reactions, we treated all samples with 10 μl of 5% sodium dodecyl sulfate (SDS) followed by 10 μl of 250 mM Na_2_EDTA (pH 8.0). We then treated the reaction mixtures with 10 μl of PK and incubated them at 45 °C for 30 min to digest the denatured topoisomerases. We extracted the DNA with phenol:chloroform:isoamyl alcohol (25:24:1) and precipitated it with 100% ice-cold ethanol. We resuspended the DNA in 20 μl of 5 mM Tris (pH 8.0) and 0.5 mM EDTA (pH 8.0) and assessed it for purity by the A260/A280 absorbance ratio.

### SHAN-seq library preparation, sequencing, and analysis

We performed SHAN-seq library preparation as previously described (15). Following the DNA cleavage reaction, we treated the DNA with TDP2 to remove the 5′ phosphotyrosine adducts and purified it with a PCR Clean-up kit (Qiagen). We added three synthetic DNA oligonucleotides with different sequences to the TDP2-treated DNA at a molar ratio of 1:100, 2:100, and 4:100 to normalize read counts across each sample after sequencing and permit accurate comparison. We then performed two rounds of NEBNext® Ultra™ II DNA PCR-free Library Prep Kit for Illumina® and NEBNext® Multiplex Oligos for Illumina® (Unique Dual Index UMI Adapters DNA Set 1) according to the manufacturer’s instructions. Each round of library prep contains an end-repair step, an A-tailing step, and an adaptor ligation step. We performed all cleanup and size selection steps with SPRI beads (Beckman Coulter).

After the first round of library preparation, we fragmented the DNA to an average length of 500 bp with an ME220 focused-ultrasonicator (Covaris). The length distribution after sonication was verified on a Tapestation (Agilent). After the second round of library preparation, we measured the concentration of adapter-labelled DNA fragments in each sample via qPCR using an NEBNext® Library Quant Kit for Illumina® (NEB). We then pooled the samples at an equimolar concentration and performed paired-end sequencing on a NovaSeq 6000 (SP flow cell, 300 million reads) to a total read length of 50 bp.

We trimmed the sequenced reads with Trimmomatic (42) and mapped them to the pBR322 plasmid reference sequences with BWA-MEM (43). We sorted and indexed the BAM files with SAMtools (44). We performed all subsequent analysis steps with custom-written Python scripts. We extracted raw plasmid read counts from BAM files using Pysam (https://github.com/pysam-developers/pysam) and normalized them by the spike-in DNA read counts. After accounting for the characteristic offset between forward and reverse reads from the staggered double-stranded breaks, we identified cleavage sites as positions with both forward and reverse read count peaks that were greater than three standard deviations above background. Then, we combined the forward and reverse read counts to get the total counts. To calculate sequence preferences, we extracted the cleavage site sequences and aligned them. We calculated the frequency of each base at each position, weighting each sequence by its read counts, and compared it to the frequency of each base occurring in the pBR322 reference sequence. We performed all statistical tests using Scipy (45) and visualizations with matplotlib (46).

## Results

### Mapping gyrase- and topoisomerase IV-mediated DNA cleavage with SHAN-seq

To characterize species-specific differences in the specificity of DNA cleavage mediated by type II topoisomerases, we mapped and compared the cleavage specificities of gyrase and topoisomerase IV from three different pathogenic bacterial species, *Ec, Ba*, and *Mt*. We previously used our SHAN-seq method to map *Ec* gyrase and topoisomerase IV cleavage on negatively supercoiled pBR322 in the presence and absence of the fluoroquinolone, ciprofloxacin (15). Therefore, to compare to these results, we performed DNA cleavage assays with *Ba* gyrase and topoisomerase IV, as well as *Mt* gyrase [the only type II topoisomerase in this species (9, 40)], on -sc pBR322 in the presence and absence of saturating ciprofloxacin (100 µM). To maintain the topological state of the DNA, we did not include ATP in our cleavage assays. Gyrases have been reported to slowly relax -sc DNA even in the absence of ATP (47), however, we did not observe appreciable -sc relaxation in any of our cleavage assays.

After performing SHAN-seq on all the samples and sequencing them, we aligned the reads onto the pBR322 plasmid reference sequence, and mapped the number of reads that aligned to each position on the plasmid (Fig. 1). In the presence of ciprofloxacin, we found a high degree of correlation between replicate read counts for each enzyme (Pearson correlation coefficient, *r* > 0.9, Supplementary Fig. 1), indicating excellent reproducibility. We also found a high degree of correlation between the forward and reverse read counts after accounting for the characteristic offset due to the staggered double-stranded break created by gyrase and topo IV cleavage (*r* > 0.9, Supplementary Fig. 1). In the absence of ciprofloxacin, we found an inconsistent or low correlation between replicate read counts (Supplementary Fig. 2), likely due to low intrinsic cleavage levels, so we did not compare these to our previous *Ec* results.

**Figure 1:**
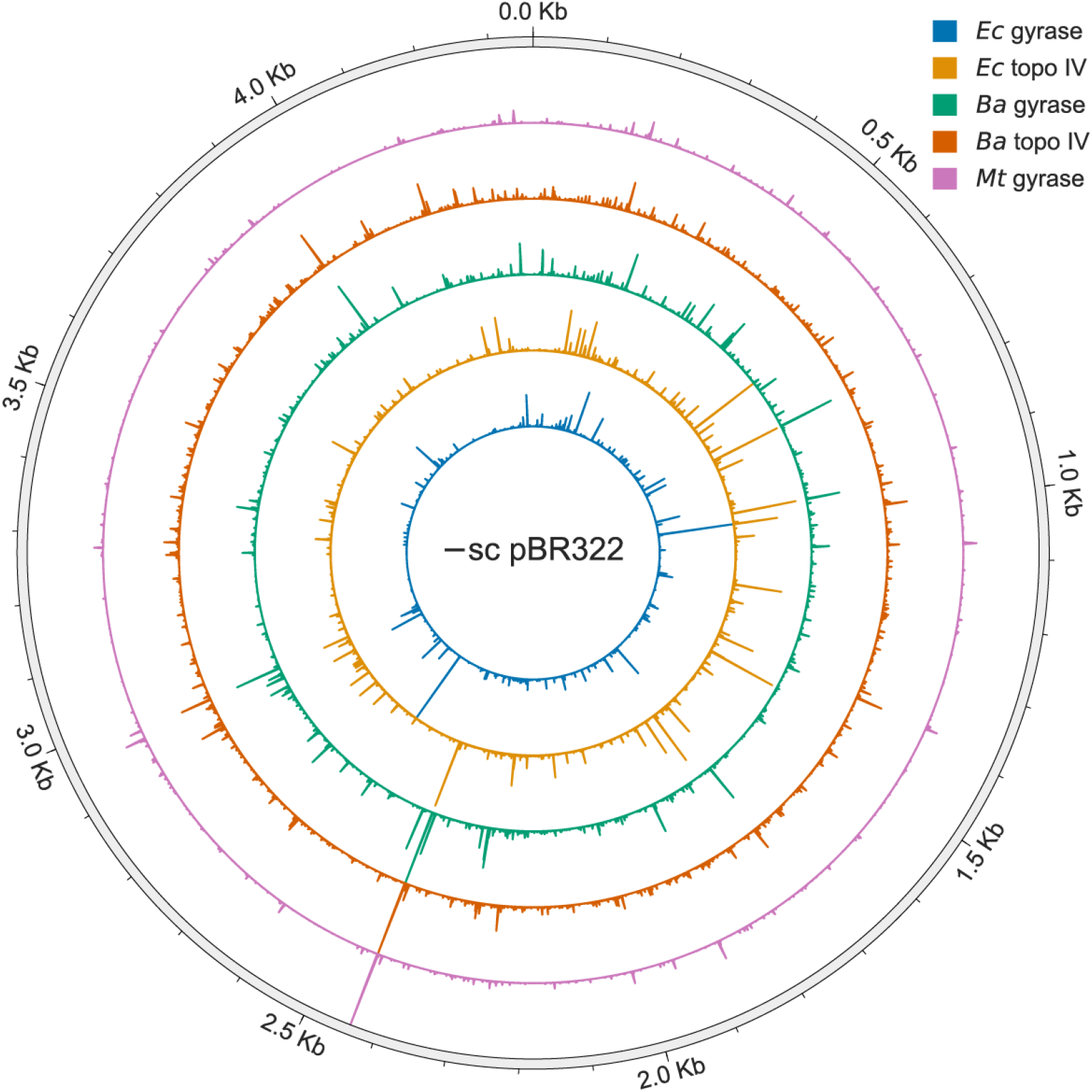
Map of DNA cleavage by *Ec* gyrase and topo IV, *Ba* gyrase and topo IV, and *Mt* gyrase on -sc pBR322 plasmid DNA in the presence of ciprofloxacin. Height of bars show the relative cleavage at each position on the plasmid DNA. The DNA cleavage maps of *Ec* gyrase and topoisomerase IV are from Ref 13.

### Gyrase and topoisomerase IV cleavage levels vary in an enzyme- and species-specific manner

First, we compared gyrase- and topoisomerase IV-mediated DNA cleavage at the well-characterized strong gyrase cleavage site at position 991 on pBR322 (Fig. 2A) (48). Previous studies suggested that gyrase and topoisomerase IV from different species have similar cleavage specificities because they all cleave this site (12), but we found that only *Ec* gyrase and topoisomerase IV have high DNA cleavage levels at the site. This result indicates that although various gyrases and topoisomerases IV may cleave the same DNA sites, they can do so with widely varying cleavage levels.

**Figure 2:**
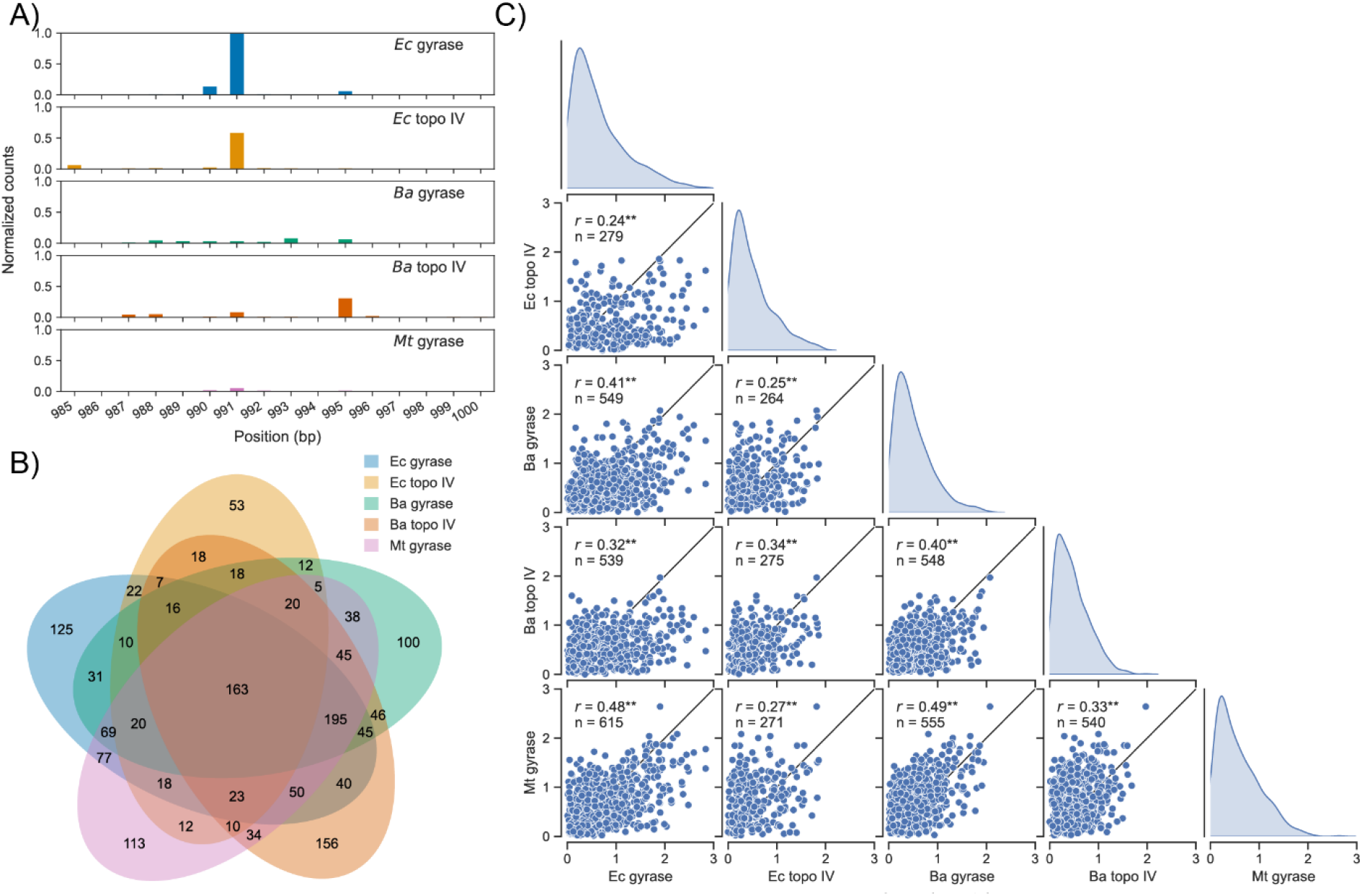
The specificity of gyrase and topo IV cleavage varies in an enzyme- and species-specific manner on -sc pBR322 plasmid DNA in the presence of ciprofloxacin. (A) Read counts around the well-characterized strong gyrase cleavage site on pBR322 at position 991. (B) Overlap in cleavage sites identified for each enzyme. (C) Pair plots showing histograms and scatter plots of log10 (read counts) for sites cleaved by two or more enzymes. Pearson correlation coefficient (*r*) and the number of DNA cleavage sites (*n*) are shown on each plot. Black lines represent *r* = 1 and ** represents *p*-value ≪ 0.005.

Next, we identified and compared all the DNA cleavage sites on the plasmid for each enzyme.

Across all the enzymes, we identified thousands of cleavage sites (2143 sites) on -sc pBR322 and hundreds of cleavage sites for each enzyme (427-911 sites). Despite the large number of DNA cleavage sites for each gyrase and topoisomerase IV, only a fraction of them were shared across all the enzymes (<37%, Fig. 2B). Furthermore, the DNA cleavage levels at sites that were cleaved by two or more enzymes were only poorly-to-moderately correlated across enzymes (*r* < 0.5, Fig. 2C). Notably, gyrase DNA cleavage levels were more correlated (*r* > = 0.4, Fig. 2C) than topoisomerase IVs (*r* < 0.4), suggesting that the DNA cleavage specificities of gyrases are more similar across species than topoisomerases IV. Additionally, between gyrase and topoisomerase IV, the *Ba* enzymes were more correlated (*r* = 0.40) than the *Ec* enzymes (*r* = 0.34), indicating that the cleavage specificity of *Ba* gyrase and topoisomerase IV are more similar than the *Ec* enzymes. Overall, these results reveal complex enzyme- and species-specific variations in cleavage specificities.

### Gyrase cleavage specificities are sensitive to DNA supercoil chirality

Although gyrases typically maintain fewer cleavage complexes on +sc than on -sc DNA (8, 9), we previously found that *Ec* gyrase and topoisomerase IV DNA cleavage specificities were relatively insensitive to supercoil chirality (15). Specifically, we found that the enzymes cleaved many of the same sites on -sc and +sc pBR322 DNA (> 85% of sites) and the cleavage levels at these sites, although universally lower on +sc DNA, were relatively well-correlated (*r* > 0.75, Fig. 3). Due to differences in cleavage specificities across gyrase and topoisomerase IV among bacterial species, we examined the effects of supercoil chirality for *Ba* gyrase and topo IV and *Mt* gyrase cleavage specificity. We performed identical cleavage and SHAN-seq analysis with the enzymes on +sc pBR322 in the presence of ciprofloxacin. For all enzymes, we found a high degree of correlation between replicate read counts (*r* > 0.99, Supplementary Fig. 1), indicating excellent reproducibility, and between forward and reverse reads after accounting for the expected offset due to the staggered double-stranded breaks from gyrase and topo IV cleavage (*r* > 0.99, Supplementary Fig. 1).

**Figure 3:**
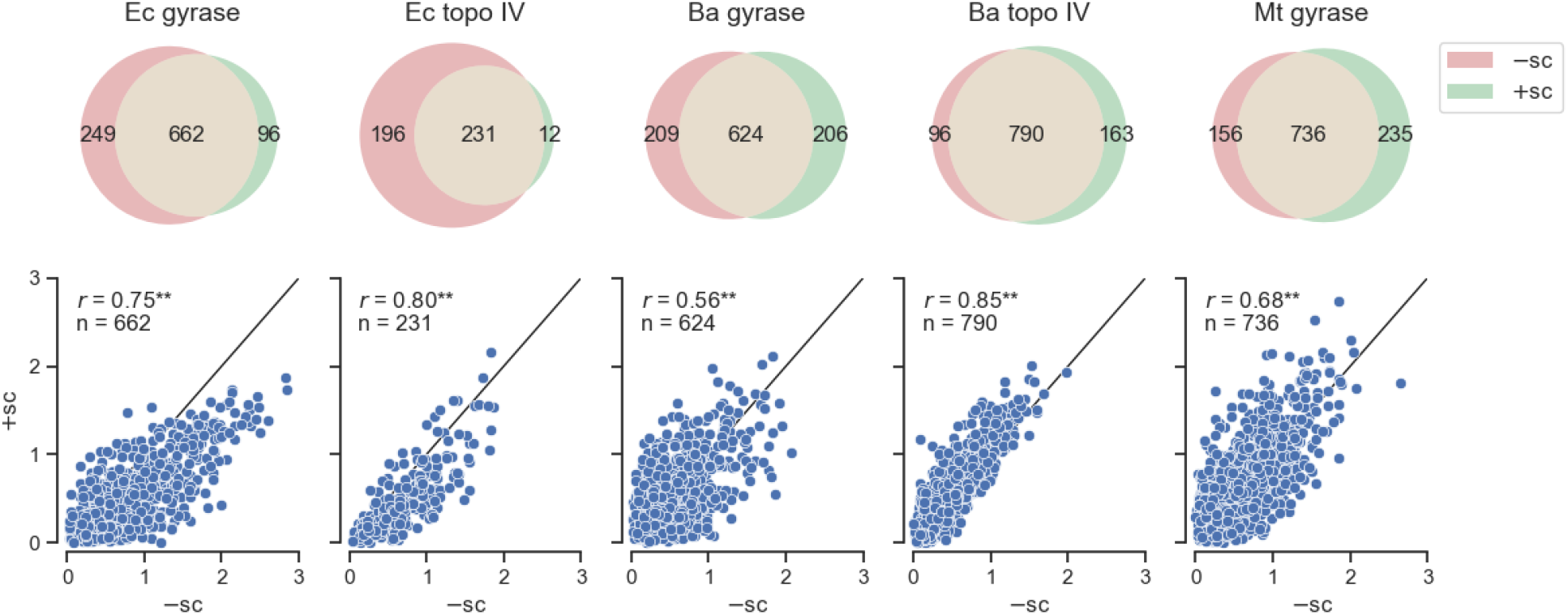
Gyrase cleavage specificities are sensitive to supercoil chirality. (Upper) Venn diagrams of the overlap in cleavage sites identified on -sc and +sc pBR22 for each enzyme in the presence of cip. (Lower) Scatterplots of the normalized cleavage levels for cleavage sites that were identified on both -sc and +sc pBR322 DNA for each enzyme in the presence of cip. Black lines represent *r* = 1 and ** represents *p*-value ≪ 0.005.

Similar to our results with -sc pBR322, we identified more than a thousand cleavage sites (1628 sites) across all enzymes and hundreds of cleavage sites for each enzyme (Fig. 3, 243-971 sites). We found similar cleavage specificity differences across the enzymes with a higher correlation between gyrase (*r* > 0.43) than topo IVs (*r* = 0.26) across species (Supplementary Fig. 3). Similar to *Ec* topo IV, we found that *Ba* topoisomerase IV cleaves many of the same sites on -sc and +sc pBR322 DNA and the cleavage levels at these sites were well-correlated (*r* = 0.85). However, when we compared the DNA cleavage sites for *Ba* and *Mt* gyrase, we found that the overlap on -sc and +sc DNA was less than 85% and the read counts at overlapping sites were less correlated than *Ec* gyrase. Overall, gyrase cleavage specificities tended to be more sensitive to supercoil chirality than topoisomerase IV cleavage specificities. These results suggest that gyrase cleavage specificities may be sensitive to supercoil chirality in some species.

### Interactions between the G-segment and the CTDs account for differences in bacterial type II topo cleavage specificities

To gain further insight into DNA cleavage specificity differences among the type II topoisomerases, we calculated and analyzed the sequence preferences for the *Ec* and *Ba* gyrase and topoisomerase IV and *Mt* gyrase. For each enzyme, we determined the sequences around the cleavage sites, aligned them, and calculated the enrichment of each base at each position, accounting for ‘weak’ and ‘strong’ sites by weighting each cleavage site sequence by its read counts. By convention, we denote the 5′/3′ nucleotides immediately adjacent to the nicks on each strand as -1/+1 and +4/+5 (Fig. 4A). Based on this analysis, we identified two regions with significant sequence preferences (Supplementary Fig. 4A), an approximately 20 bp region with strong biases [-log(p) > 10)] surrounding the cleavage site that corresponds to the DNA bound by the cleavage domain and an extended 120 nucleotide region with weak biases (−log(p)>3) that corresponds to the DNA interacting with the CTDs of the enzymes (Fig. 4A). Both regions were palindromic in their sequence preferences with respect to the center of the cleavage site.

**Figure 4:**
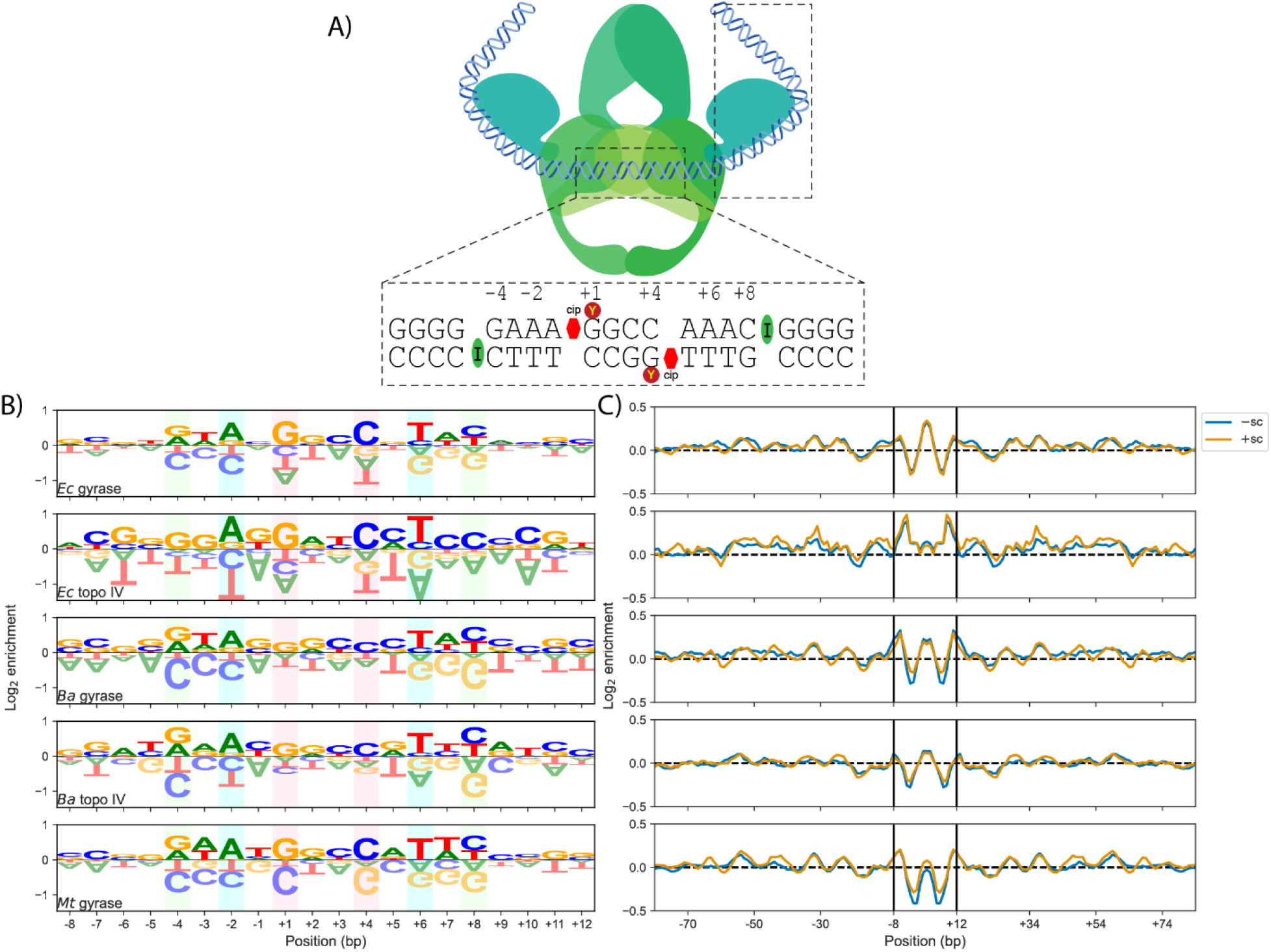
Gyrase and topo IV cleavage site sequence preferences. (A) Diagram of protein regions that interact with DNA during cleavage. The catalytic core interacts with approximately 20 bp surrounding the cleavage site, whereas the CTDs interact with up to 120 bp surrounding the cleavage site. Within the catalytic core, an isoleucine (I) residue intercalates between bases -5/-4 and +8/+9, a tyrosine (Y) residue covalently interacts with the +1 and +4 bases, and ciprofloxacin inserts between the -1/+1 and +4/+5 bases. (B) DNA Cleavage site sequence preferences of the catalytic core region and (C) GC preferences with a five bp rolling mean including the CTD regions for each enzyme. Enrichment represents the frequency of a base in the cleavage site over the background frequency of that base in pBR322. The green, blue, and red boxes highlight the most commonly enriched bases.

In agreement with previous results (12, 15, 22, 23), the most significant sequence preferences within the cleavage domain region were -4G/+8C, -2A/+6T, and +1G/+4C (Fig. 4B). However, whereas the -4G/+8C and -2A/+6T biases were consistent across all of the enzymes, the +1G/+4C bias varied considerably across the enzymes in a species-specific manner. Additionally, there were minor sequence preference differences between enzymes and across species, particularly at the -3/+7 and -1/+5 positions. Given that the +1G/+4C preference is associated with the intercalation of quinolones (13, 15), these results suggest that sequence or structural differences across the enzymes affect fluoroquinolone stabilization.

In addition to the sequence preference variations within the cleavage domain, there were differences in the sequence preferences associated with the CTD region (Fig. 4C). Most notably, all of the gyrases had periodic GC preferences with a 10-11 bp periodicity (Supplementary Fig. 4B), indicative of their ability to wrap DNA around their CTDs (22). We also observed a similar, but much weaker, periodic GC preference for *Ba* topo IV. The periodic GC preferences were much more pronounced on +sc than on -sc DNA, particularly for *Ba* and *Mt* gyrase, suggesting that the sensitivity of the enzymes to DNA supercoil chirality is due in part to the interaction of their CTDs with DNA. More broadly, our results show a number of subtle differences within the cleavage domain and CTD regions that likely contribute to the significant differences among the cleavage specificities of individual enzymes.

## Discussion

As the physiological importance of the DNA cleavage specificity for bacterial type II topoisomerases continues to emerge, there is considerable interest in the factors that control the cleavage-religation equilibrium (26, 29). Previous work has shown that variations in sequence and structure of gyrases and topoisomerase IV within and across bacterial species can influence their catalytic activities, as well as their susceptibility to fluoroquinolones. However, it is unclear whether these variations also affect DNA cleavage specificity (2, 26, 29, 30). Using SHAN-seq, we found that the DNA cleavage specificity of gyrases and topoisomerase IV differ within species, across bacterial species, with supercoil chirality, and in response to ciprofloxacin. Our results suggest that bacteria tune the specificity of gyrase and topoisomerase IV cleavage, which may, in turn, affect their susceptibility to fluoroquinolone antibacterials.

To resolve DNA entanglements throughout the bacterial genome, gyrase and topoisomerase IV need to be able to cleave DNA wherever these entanglements occur. Consistent with our previous observations and the physiological requirement of abundant cleavage sites, we identified hundreds of cleavage sites for each enzyme on pBR322, more than one site every ten bp (Fig. 1 and 2). Despite the large number of DNA cleavage sites, we found that scission levels at these sites varied by as much as two-to-three orders of magnitude (Fig. 2C). Such a broad variation in cleavage levels highlights the strong regulatory effect of DNA sequence on the cleavage-religation equilibrium. These broad variations likely represent differences in the enzymes’ ability to bind, bend, cleave, and/or religate different DNA sequences. Unfortunately, the equilibrium nature of our experiments precludes us from being able to differentiate among these factors. Single-molecule studies on yeast topo II found that the ability of the enzyme to bend different DNA sequences determined its cleavage levels (49), but further studies are needed to determine whether gyrases and topoisomerase IV behave similarly.

We found that type II topoisomerases from different bacterial species varied considerably in their DNA cleavage site preferences (Fig. 2). This suggests that DNA sequence has a strong regulatory effect on the enzymes’ cleavage-religation equilibria. Across all the enzymes, the DNA bound at the cleavage domain had some similar sequence preferences, particularly at the -4G/+8C and -2A/+6T, and +1G/+4C positions (Fig. 4B). Previous structural work has shown that a conserved isoleucine residue inserts between the -5/-4 and +8/+9 bases, accounting for the -4G/+8C preference (13, 50, 51). Similarly, previous structural work has shown that fluoroquinolones insert between the -1/+1 and +4/+5 bases and interact with the bases via base-stacking or pi-pi interactions, accounting for the +1G/+4C (Fig. 4A). The origin of the -2A/+6T preference is unknown, but has previously been observed both in the presence and absence of fluoroquinolones (13, 15, 19, 23, 50, 51). However, between gyrases and topoisomerase IVs and across species, there were several notable sequence preference differences.

The largest difference between the sequence preferences of the gyrases and topoisomerases IV examined in the current study occurred beyond the region bound by the cleavage domain (Fig. 4C). All of the gyrases had strong extended periodic GC sequence preferences within 120 bp of their cleavage sites that are associated with the abilities of their CTDs to wrap DNA (15, 22). This finding is in contrast to those of the topoisomerases IV, which do not wrap DNA. Unexpectedly, *Ba* topoisomerase IV displayed a weak periodic GC sequence preference similar to gyrase (Fig. 4C and Supplementary Fig. 4), which likely accounts for its cleavage specificity being better correlated with *Ba* gyrase than *Ec* topoisomerase IV was correlated with *Ec* gyrase (Fig. 2C). Although the CTD of *Ba* topoisomerase IV lacks the conserved GyrA box motif found in gyrases, it shares the six blade structure with the gyrases CTDs as opposed to the five blade structure of the *Ec* topo IV CTD (Supplemental Fig. 5). This six-bladed CTD structure may enable the CTD of *Ba* topoisomerase IV to interact with DNA in a manner akin to gyrases. Together, our results suggest that the differences in the CTDs of gyrases and topoisomerase IV are likely responsible for variations in cleavage specificity within species.

We also found significant differences in the DNA cleavage specificity of gyrases and topoisomerases IV across species. Gyrases tended to be better correlated with other gyrases across species than topoisomerases IV, likely due to similarities in the sequence and structure of their CTDs. However, they were still poorly correlated overall (*r* < 0.5, Fig. 2C). Within the cleavage domain region, there were mostly minor sequence preference variations across the enzymes, particularly at the -3/+7 and -1/+5 positions (Fig. 4B). All the enzymes showed the -4G/+8C, -2A/+6T, and +1G/+4C preferences. Yet, whereas the magnitude of the -4G/+8C and -2A/+6T preferences was similar across all the enzymes, the magnitude of the +1G/+4C preference varied considerably in a species-specific manner. Specifically, the *Ec* and *Mt* type II topoisomerases had a much greater preference for +1G/+4C than the *Ba* type II topoisomerases. The +1G/+4C preference is thought to be associated with base-stacking and pi-pi interactions between fluoroquinolones and guanine bases (52). This result could be explained if *Ba* gyrase and topoisomerase IV DNA cleavage complexes were not saturated with ciprofloxacin. However, we used saturating levels of ciprofloxacin (100 µM) in all our experiments (9). One possibility is that *Ba* gyrase and topoisomerase IV position ciprofloxacin within their cleavage complexes in an orientation that disfavors base-stacking interactions. Although some structural studies have speculated that base-stacking interactions between fluoroquinolones and topoisomerase-cleaved DNA could differ across gyrases and topoisomerases IV from different bacterial species and various fluoroquinolones, base-stacking interactions are notoriously difficult to estimate from structures (36). Notably, we previously showed that ciprofloxacin-stabilized *Ba* gyrase and topoisomerase IV cleavage complexes persist for much longer than *Ec* gyrase and topoisomerase IV or *Mt* gyrase cleavage complexes (9). Hence, base-stacking interactions with fluoroquinolones may be more important when the topoisomerase-fluroquinolone interactions are less stable. More broadly, our results suggest that minor variations in topoisomerase sequence and structure that have subtle effects on the enzymes’ DNA sequence preferences may have much larger effects on their susceptibility to fluoroquinolones.

Previously, we found that the specificity of DNA cleavage mediated by *Ec* gyrase and topoisomerase IV was insensitive to supercoil chirality (15). Yet, given the differences within and across species, we decided to examine the effects of supercoil chirality on the cleavage specificity of each enzyme. While the specificity of *Ba* topoisomerase IV cleavage was relatively insensitive to supercoil chirality, we unexpectedly found that the specificity of *Ba* gyrase and *Mt* gyrase cleavage was sensitive to supercoil chirality. In analyzing their sequence preferences, we found that these enzymes tended to have weaker periodic GC sequence preferences on -sc DNA than +sc DNA (Fig. 4C), suggesting that the difference in sensitivity to supercoil chirality is largely due to variations in the ability of CTDs to interact with DNA. Previous work has shown that *Mt* gyrase and *Bacillus subtilis* gyrase, which is closely related *Ba* gyrase, have CTDs with a shorter C-terminal tail than *Ec* gyrase that reduces their ability to wrap and negatively supercoil DNA (29, 39). Thus, we attribute the sensitivity to supercoil chirality of *Mt* and *Ba* gyrases cleavage specificities to their shorter C-terminal tails and reduced ability to wrap -sc DNA.

Bacteria vary widely in their metabolism, genome content, and genome size. For example, the percent GC contents of *Mt, Ec*, and *Ba* are approximately 35, 50, and 60%, respectively (53–55). Our results strongly indicate that the cleavage specificity of gyrases and topoisomerases IV vary in a species-specific manner due to variations in the sequence and/or structure of their cleavage domains and CTDs. We posit that the variations in cleavage specificities help target enzyme activity to specific regions of the genome with DNA entanglements. Consequently, because fluoroquinolones prevent relegation mediated by type II topoisomerases in a DNA sequence-dependent manner, variations in their sequence preferences could alter the susceptibility of enzymes to fluoroquinolones. Therefore, it may be helpful to consider the specificity of gyrases and topoisomerase IV cleavage when developing new antibacterials that target these enzymes.

## Acknowledgements

This work was supported by the Intramural Research Program of the National Heart, Lung, and Blood Institute (NHLBI) of the National Institutes of Health (NIH) [1ZIAHL001056 to K.C.N.]; a National Institute of General Medical Sciences (NIGMS) fellowship [1FI2GM142536 to I.L.M.]; and NIH grants [R01 GM126363 and R01 AI170546 to N.O.]. Funding for open access charge was provided by NHLBI intramural funding to K.C.N. (ZIAHL001056). This work utilized the resources of the NHLBI DNA Sequencing and Genomics Core and NIH HPC Biowulf cluster (http://hpc.nih.gov). The contributions of the NIH author(s) were made as part of their official duties as NIH federal employees, are in compliance with agency policy requirements, and are considered Works of the United States Government. However, the findings and conclusions presented in this paper are those of the author(s) and do not necessarily reflect the views of the NIH or the U.S. Department of Health and Human Services.

## Data availability

All sequencing data are available from the NCBI GEO database (GSE303783 and GSE267149).

