## Supplemental Information for "Bacterial type II topoisomerases cleave DNA in a species-specific manner"

### Supplementary Information for Bacterial type II topoisomerases cleave DNA in an enzyme- and species-specific manner

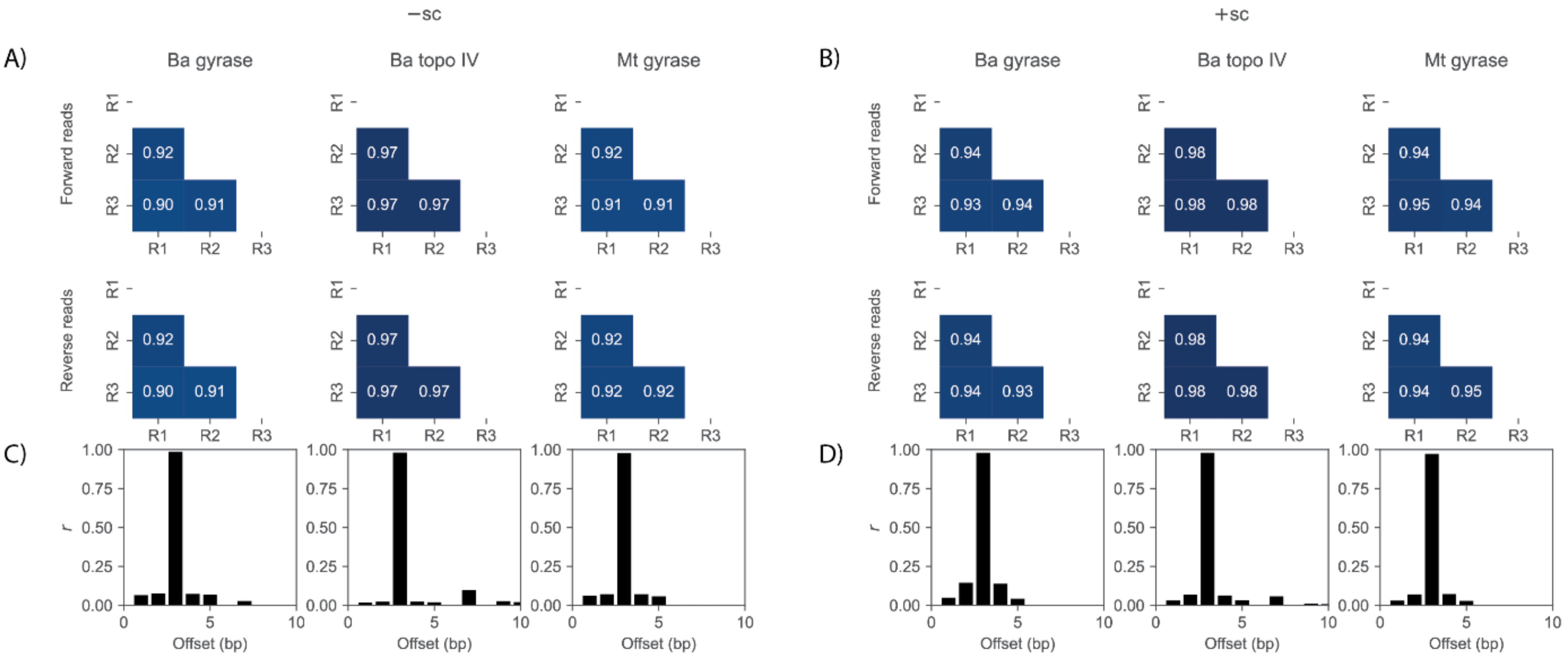


**Supplementary Figure 1:** In the presence of ciprofloxacin, cleavage by *Ba* gyrase, *Ba* topoisomerase IV (topo IV), and *Mt* gyrase produced forward and reverse read counts that were well-correlated (A) on -sc and (B) +sc pBR322 plasmid DNA, indicating excellent reproducibility. After accounting for the characteristic offset, the forward and reverse read counts were also well-correlated (C) on -sc and (D) +sc pBR322 plasmid DNA. Correlations were measured using the Pearson correlation coefficient (*r*).

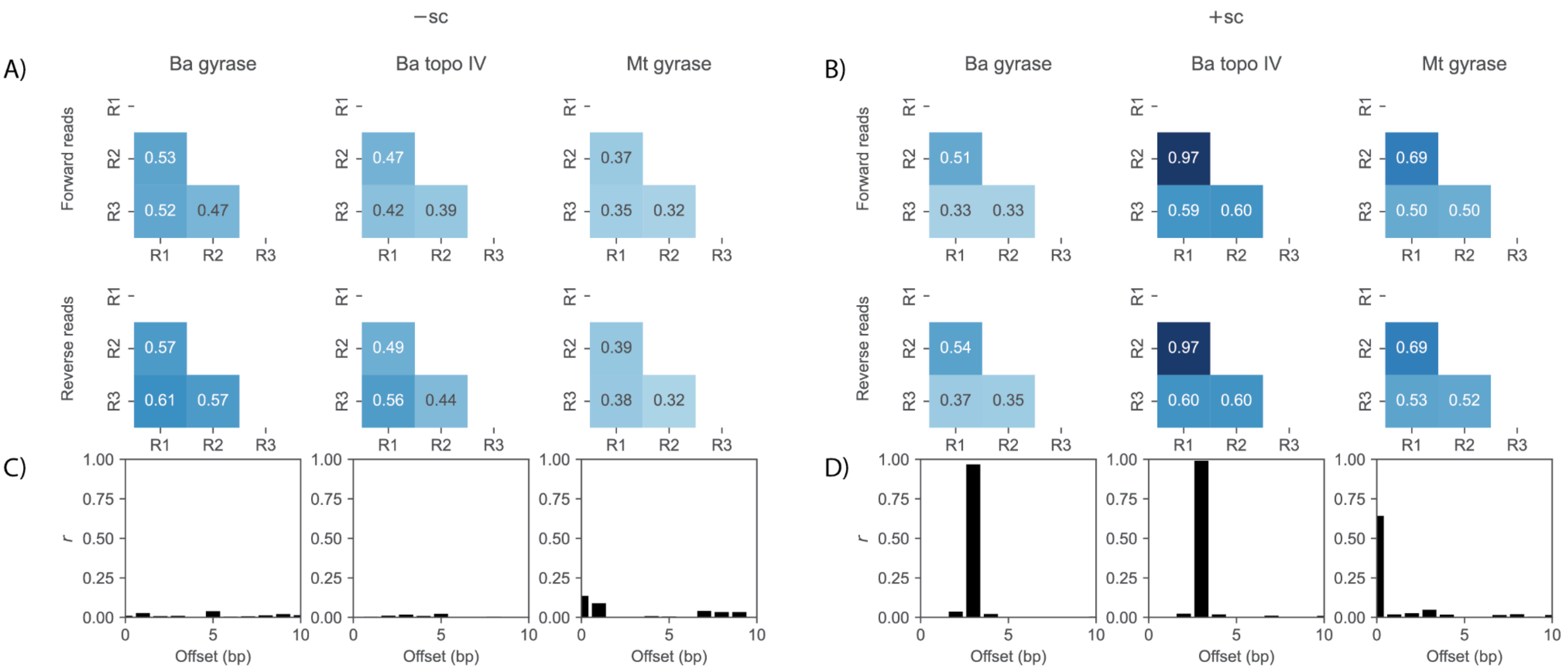


**Supplementary Figure 2:** In the absence of ciprofloxacin, cleavage by *Ba* gyrase, *Ba* topoisomerase IV (topo IV), and *Mt* gyrase produced forward and reverse read counts that were poorly to moderately correlated on (A) -sc and (B) +sc pBR322 plasmid DNA. After accounting for the characteristic offset, the forward and reverse read counts were not consistently well-correlated on (C) -sc and (D) +sc pBR322 plasmid DNA. Correlations were measured using the Pearson correlation coefficient (*r*).

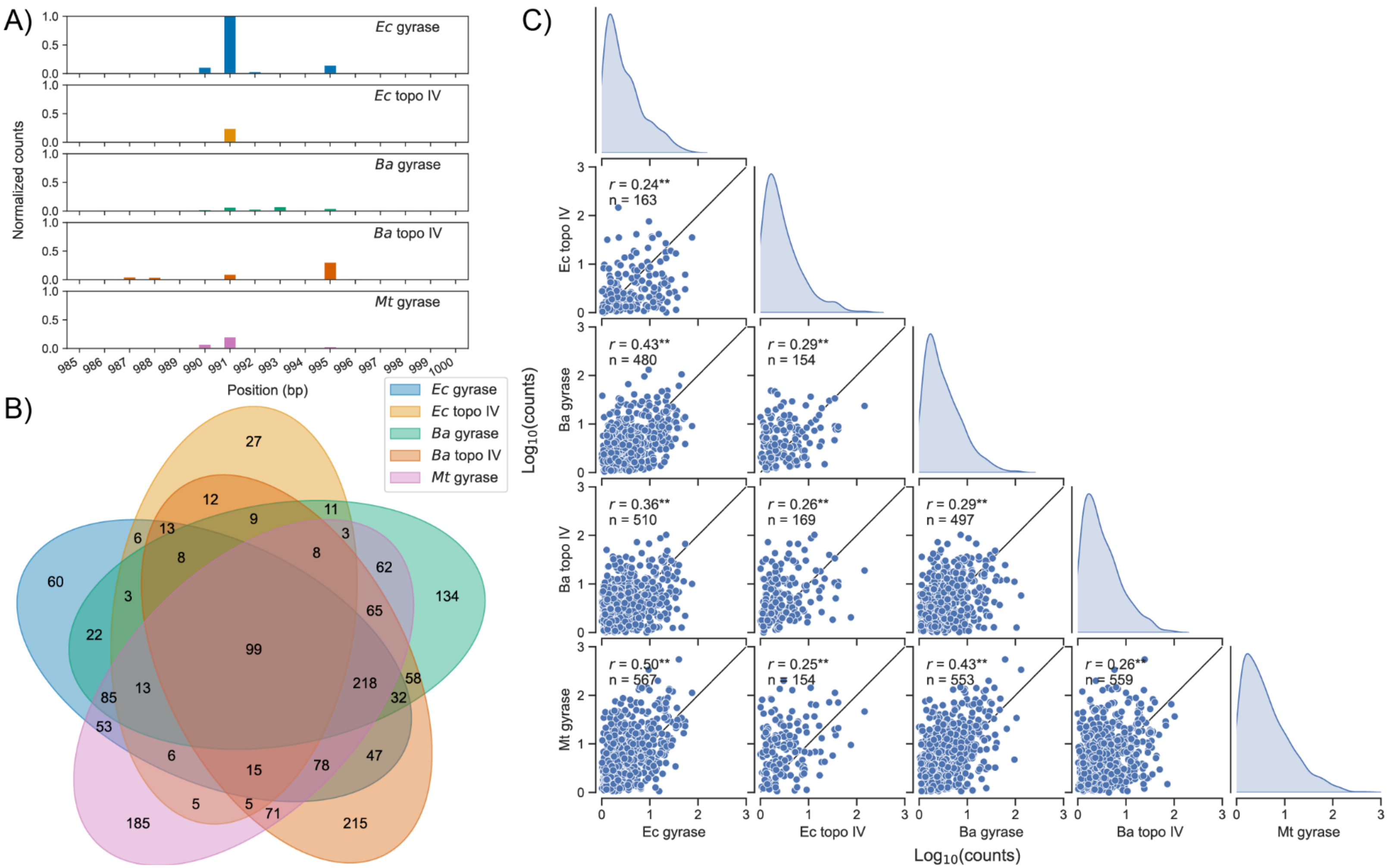


**Supplementary Figure 3:** The specificity of DNA cleavage mediated by gyrases and topoisomerases IV varies in an enzyme- and species-specific manner on +sc pBR322 in the presence of cip. (A) Read counts around the well-characterized strong gyrase cleavage site on pBR322. (B) Overlap in cleavage sites identified for each enzyme. (C) Pair plot showing histograms and scatter plots of $\log_{10}$(read counts) for sites cleaved by two or more enzymes. Pearson correlation coefficient ($r$) and the number of cleavage sites ($n$) are denoted on each plot. Black lines represent $r = 1$ and ** represents $p$-value $\ll 0.005$.

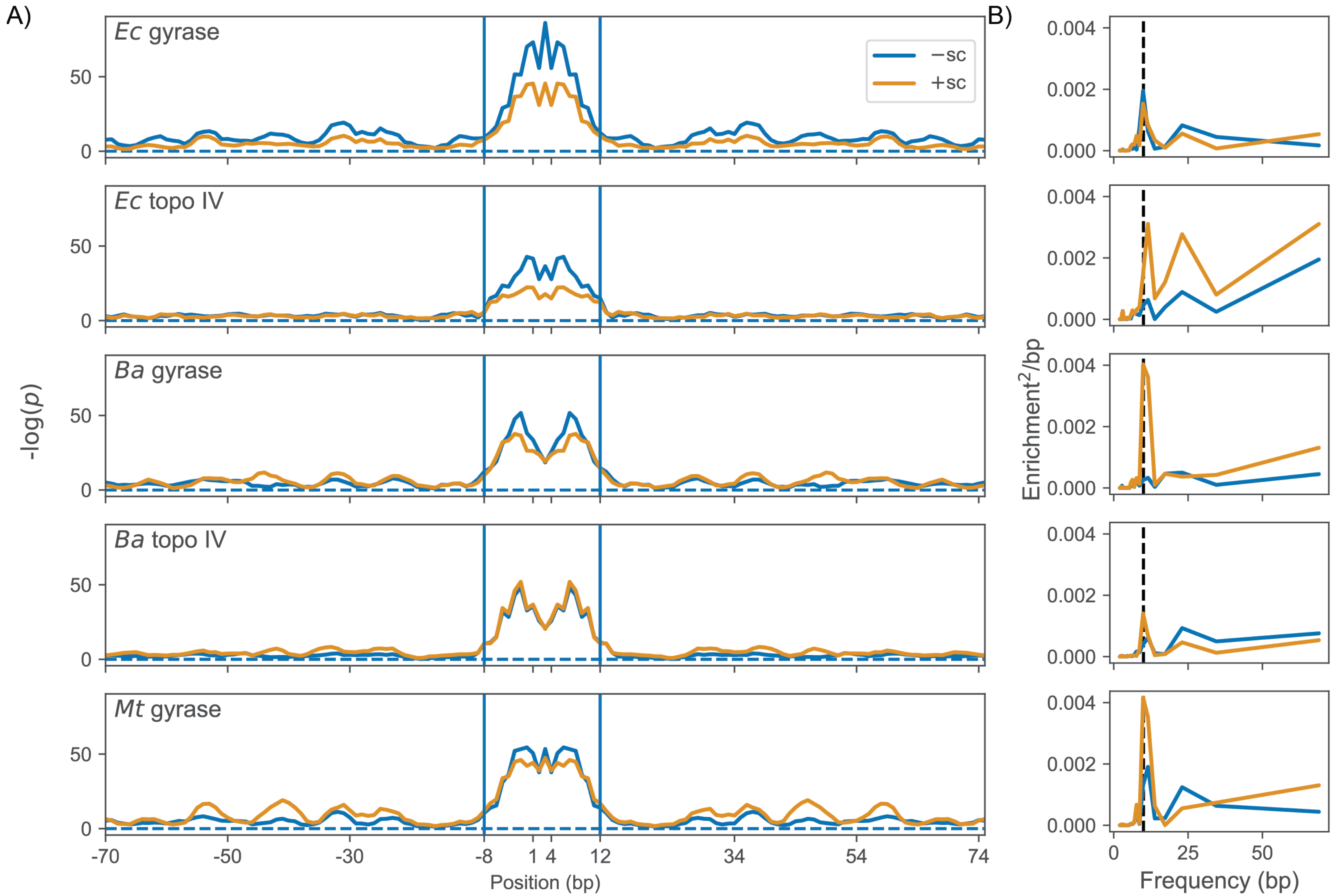


**Supplementary Figure 4:** (A) Chi-squared analysis of sequence preference biases with a 4 bp moving average on -sc and +sc pBR322 DNA. The -$\log_{10}$ of the p-value is shown. The dotted horizontal line denotes $p > 0.01$. The central region between the vertical lines is the DNA bound by the topo cleavage domain. The sequence preferences outside of the vertical lines represents the region that can interact with the CTDs. (B) Power spectral density estimation of the sequence preferences outside of the cleavage domain shows an approximately 10 bp (vertical black line) periodicity.

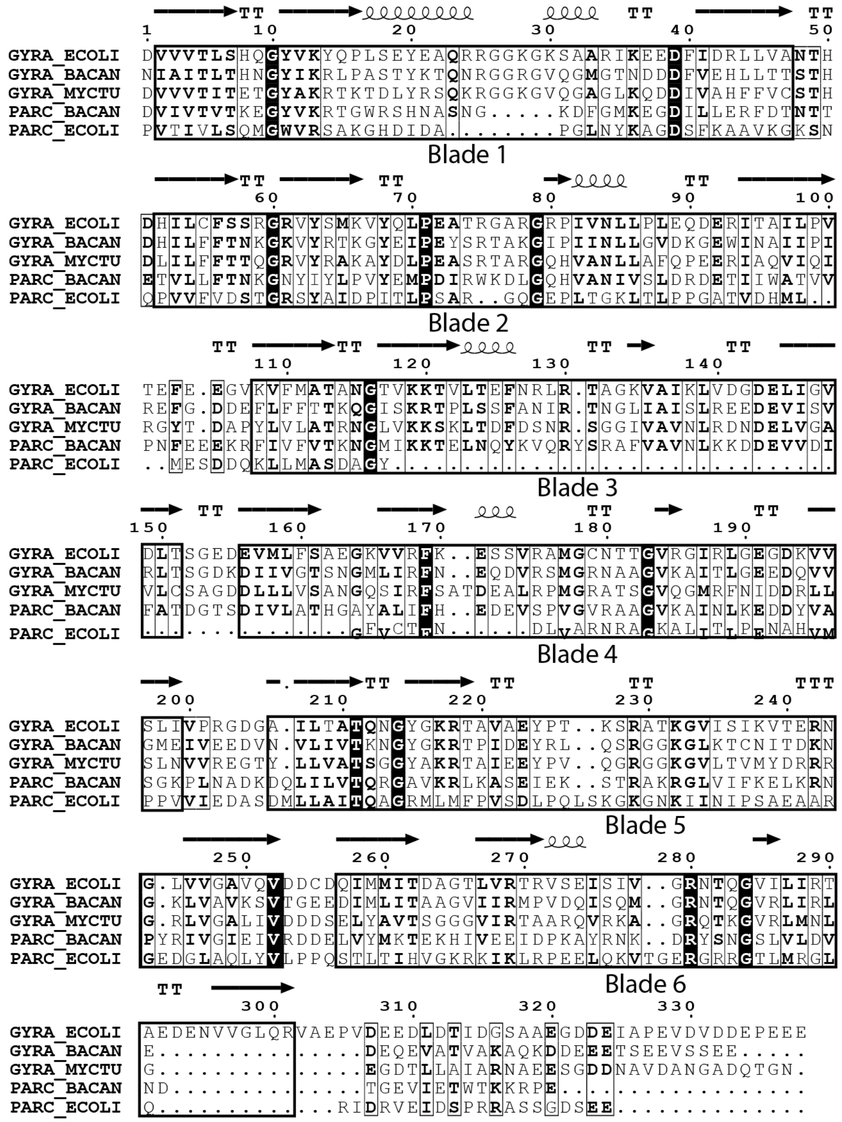

**Supplementary Figure 5:** Multiple sequence alignment of gyrase and topoisomerase IV CTDs using ClustalW with blades highlighted by black boxes. All the gyrases have six blades in their CTD but have different C-terminal tails. *Ba* topoisomerase IV has six blades, similar to the gyrases, whereas *Ec* topoisomerase IV has five blades.
